# 2D MXene-DNA Hybrid Hydrogel for Thrombin Detection: A Versatile Approach for Biomedical Sensing

**DOI:** 10.1101/2024.04.24.590924

**Authors:** Vinod Morya, Dhiraj Bhatia, Chinmay Ghoroi

**Affiliations:** Biological Engineering, Indian Institute of Technology Gandhinagar, India; Chemical Engineering, Indian Institute of Technology Gandhinagar, India

**Keywords:** DNA hydrogel, MXene, MXene hydrogel, Aptamer, Biosensing, thrombin

## Abstract

The delaminated MXene (2D MXene) and DNA hydrogel created enormous opportunities due to their versatility and ability to be tailored for specific applications. MXene offers high aspect ratio morphology and electrical conductivity, while DNA provides stimuli responsiveness and specificity in binding to ligands or complementary sequences. This synergy makes DNA an ideal actuator when combined with 2D MXenes. Present work makes the first effort to combine and exploit them for detecting the thrombin levels; a crucial proteolytic enzyme that plays a pivotal role in regulating blood clotting by cleaving fibrinogen into fibrin and plays a critical role in bleeding disorders such as Haemophilia and Von Willebrand disease. This study introduces a novel hybrid DNA hydrogel by leveraging the properties of 2D MXene with a thiol-modified thrombin binding aptamer (TBA) as a crosslinking agent. The TBA and its complementary DNA oligos are immobilized on 2D MXene sheets, forming a packed hydrogel. Upon thrombin binding, the TBA releases its complementary DNA, resulting in a loosened hydrogel and a change in resistance, which is used as a read-out for thrombin detection. The concept is successfully demonstrated, achieving 90-92% accuracy in detecting thrombin in artificial samples. This robust technique holds promise for biomedical sensing devices, allowing customization for the detection of various target molecules using specific aptamers.

## Introduction

DNA based hydrogels has emerged as a powerful tool in the field of biosensing, enabling the construction of various devices and systems by leveraging the unique properties of DNA^1^. The remarkable stability, programmability, and self-assembling properties inherent in DNA are harnessed in the development of hydrogels, thus establishing DNA as a versatile building block at the nanoscale^2^. Functional DNA motifs, including i-motifs^3^, A-motifs^4^, and aptamers^5^, play a pivotal role as intelligent actuators within stimuli-responsive hydrogels. These motifs offer unparalleled tunability, allowing for customization according to specific applications, and their potential scope is virtually limitless^6^. Among other DNA motifs, aptamers have emerged as highly valuable recognition elements in the development of sensors, owing to their outstanding affinity and selectivity towards target molecules^7,8^. Their ability to bind specifically to a desired target enables aptamers to act as effective molecular switch or trigger in a responsive systems^9^. Aptamers are short oligonucleotide sequences typically ranging from 20 to 60 nucleotides, and their specific sequences are identified through a sophisticated process called SELEX (Systematic Evolution of Ligands by Exponential Enrichment)^10^. During SELEX, a high-affinity DNA sequence is selected from a diverse DNA library through iterative rounds of enrichment for a specific ligand or target molecule. In recent years, the utilization of aptamers as sensing molecules has experienced a significant surge^11^. This is primarily attributed to their remarkable sensitivity and lower cost compared to antibodies, making them highly attractive for various sensing applications^8^. However, the organic nature of DNA imposes limitations on its physical properties, which can restrict its applicability in translating sensing readouts into tangible forms. Therefore, hybrid DNA hydrogels have been developed by incorporating DNA into various composites. This approach enables the enhancement of DNA hydrogel properties and expands their range of applications. Hybrid DNA hydrogels with an inorganic component could expands the capabilities of DNA to further level. DNA hydrogels have already been combined with a wide range of inorganic materials, including metallic nanoparticles^12^, quantum dots^13^, 2D nanosheets^14^, graphene^15^, and carbon nanotubes (CNTs)^16^, and more. These materials provide extra dimension of functionality to the DNA hydrogel.

In this work, we have successfully created a novel hybrid DNA hydrogel by combining DNA aptamer with MXene (Ti3C2Tx), a revolutionary 2D material. MXenes possess high aspect ratio morphology, high metallic conductivity and hydrophilicity^17^. We have ingeniously utilized DNA aptamer as a crosslinker to facilitate the polymerization of 2D MXene sheets, resulting in a sophisticated and intricate 3D network. Here, MXene plays a crucial role in providing electrical conductivity, while the aptamer acts as the target sensing moiety. As a proof of concept, we have designed the hybrid hydrogel to effectively detect thrombin levels in blood serum samples (Schematic 1) by using thrombin binding aptamer (TBA). The maintenance of normal haemostasis is contingent upon a well-regulated system comprising both procoagulant and anticoagulant pathways. Any disruption to these intricate processes can result in the loss of haemostatic control, leading to the risk of either excessive bleeding or thrombosis^18^. Thrombin levels play a crucial role in bleeding disorders such as Haemophilia and Von Willebrand disease. These conditions are significantly affected by the presence of inadequate or dysfunctional thrombin, leading to impaired blood clotting and increased bleeding tendencies in affected individuals^19^. Thrombin, a crucial proteolytic enzyme, plays a pivotal role in regulating blood clotting by cleaving fibrinogen into fibrin^20^. The commercially available methods to determine the blood levels are ELISA (enzyme-linked immunosorbent assay) based^21,22^. ELISA-based methods involve utilizing primary and secondary antibodies, thus incurs high costs and necessitating skilled personnel for test execution. The hydrogel’s unique properties enable it to serve as a cost effective, sensitive, and efficient platform for thrombin detection, holding great promise for potential applications in monitoring blood coagulation processes and related medical diagnostics. This system is highly configurable, allows for a wide range of potential applications in sensing and other fields.

**Schematic 1.**
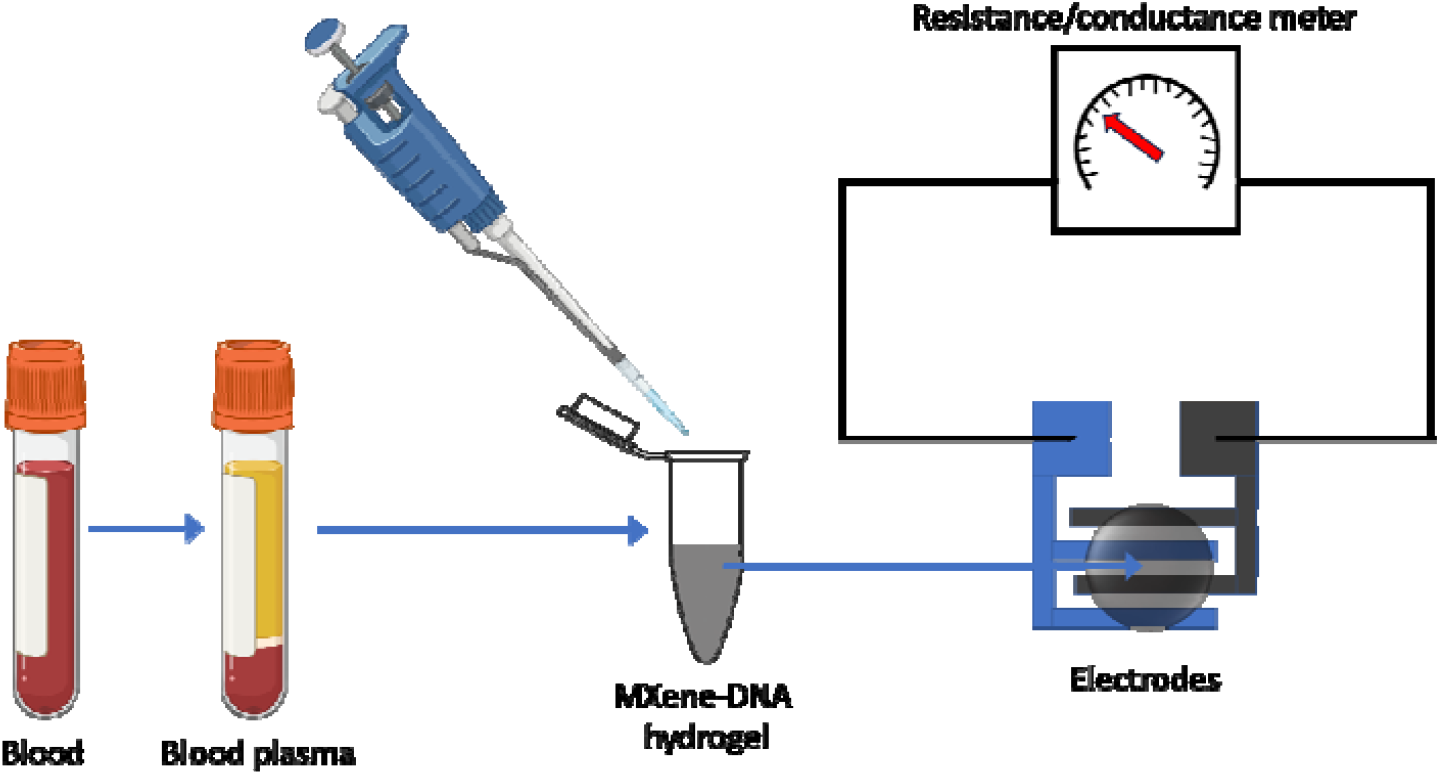
Proposed thrombin detection set-up with MXene-DNA hybrid hydrogel.

## Experimental section

### 2.1. Materials

All oligonucleotides (Table S1) at a 0.2 μM synthesis scale with desalting purification (HPLC purification in case of labelled ones) were purchased from Sigma-Aldrich (Merck). Thrombin (source-Human Plasma), Titanium aluminium carbide 312 (MAX phase) and hydrofluoric acid were also purchased from Sigma-Aldrich (Merck). Bovine serum albumin (BSA), Acrylamide/Bis-acrylamide (29:1), Ethidium bromide (EtBr), tetramethyl ethylenediamine (TEMED), paraformaldehyde and ammonium persulfate, Tris-acetate-EDTA (TAE) were purchased from HiMedia. Other salts and acid like MgCl_2_, NaCl, KCl, HCl, NaOH, CH_3_COONa, CH_3_COOH, Na_2_HPO_4_, and KH_2_PO_4_ were purchased from Finar chemicals.

### 2.2. Synthesis of MXene (Ti_3_C_2_T_x_) and delamination into 2D sheets

The MXene was synthesized using a well-documented hydrofluoric acid (HF) etching method^17^. To start, 0.5 grams of Ti_3_AlC_2_ powder was gradually added to 10 ml of 30% HF and continuously stirred for 5 hours at room temperature. Following this, the mixture was washed with deionized (DI) water using a centrifuge at 3500 rpm until a pH of 6 was reached. The resulting sediments were then resuspended and vacuum filtered using a 0.45 μm MCE (mixed cellulose ester) filter. The obtained MXene slurry was subsequently dried in a vacuum oven at 80°C for 24 hours.

For the delamination of MXene into separated 2D sheets, 0.2 grams of MXene were dispersed in Dimethyl Sulfoxide (DMSO) and left to shake overnight. During this process, DMSO infiltrates between the stacked layers of MXene, facilitating separation through sonication. The MXene was then washed 3-4 times with DI water to remove any excess DMSO. The resulting sediment was resuspended in DI water and sonicated for 6 hours to obtain a colloidal solution of delaminated MXene sheets (2D MXene).

### 2.3. Immobilization of DNA aptamer and its cDNA on amine functionalize 2D sheets

0.1 grams of vacuum-dried 2D MXene were dispersed in 10 ml of ethanol and sonicated for 30 minutes, to ensure a homogeneous dispersion. Following this, 10% APTES was added to the solution and shaken for 6 hours^23^. The solution was then washed 4-5 times with ethanol to remove any excess APTES, and the resulting pellet was vacuum dried.

For DNA immobilization, 5 mg of APTES-functionalized 2D MXene was dispersed in 1 ml of PBS (pH 7.2) and sonicated for 5 minutes. Next, 10 μl of Sulfo-MBS solution was added and shaken for 30 minutes at room temperature (RT). The mixture was then centrifuged at 10,000 rpm to remove any excess Sulfo-MBS. The resulting pellet was dispersed in 200 μl of PBS, and 10 μl of each Apt1 and Apt2 were added, followed by incubation for 30 minutes at RT. The solution was centrifuged again at 10,000 rpm to remove any unbound oligos, and the pellet was redispersed in PBS and stored at 4 °C.

### 2.4. Characterization

#### Electrophoretic Mobility Shift Assay (EMSA)

For EMSA, 10% native polyacrylamide gel electrophoresis (PAGE) was used. For sample loading, the solution was diluted up to 5 μM with buffer (1X TAE) solution then mixed with 1X loading dye and kept it for 3 min to allow the dye integration with the DNA completely. The samples were then loaded into the wells and run at 10 volts cm-1 for 80 min in 1× TAE running buffer. Finally, the gels were stained with ethidium bromide (EtBr), and then scanned using a ‘Bio-Rad ChemiDoc MP’ imaging system.

#### UV-Visible (UV-Vis) spectroscopy

Absorption studies were conducted using the ‘Analytik Jena Specord 210 Plus’ UV-visible spectrophotometer with a quartz cuvette having a working volume of 1 ml. This instrument allowed us to analyze the absorbance properties of the samples across the UV-visible spectrum, providing valuable insights into the conversion and functionalization of MXene.

#### Fourier transform infrared (FT-IR) spectroscopy

FTIR spectra were recorded in the range of 4000-450 cm-1 using ‘Perkin Elmer spectrum two’ FT-IR Spectrophotometer.

#### Dynamic Light Scattering (DLS)

The reduction in size during the conversion from MXene to 2D MXene sheets was assessed by monitoring the changes in their hydrodynamic size. A quartz cuvette of 1 mL working volume is used with ‘Malvern Panalytical Zetasizer Nano ZS’ instrument.

#### Field Emission Scanning Electron Microscope (FE-SEM)

The morphology of the different stages of MXene was analysed using a ‘JEOL, JSM-7900F’ FE-SEM instruments. Powder samples of MAX phase and MXene were sprinkled directly on carbon tap. Other samples were prepared by drop casting on to silicon wafer substrate. Prior to imaging, platinum was coated on the sample through sputtering for 90 s to make the surface conductive.

#### Atomic Force Microscope

The morphological characterization of the MXene-DNA hybrid hydrogel was performed on ‘MFP-3D Bio-AFM’ (Asylum Research, Oxford Instruments) with AC160TS (Olympus) cantilever in tapping mode. For the imaging, 10-20 μL of samples were drop-casted on an acid washed glass substrate and dried in vacuum oven. AFM scanning was performed with the peak force tapping mode to analyse the surface morphology.

### 2.5. Conductivity measurements

Platinum (Pt) was deposited onto an acid-cleaned glass substrate using a sputtering technique (refer to Figure S1). The substrate preparation involved cutting glass slides into square shapes (∼25 mm) and sequentially washing them with concentrated nitric acid, acetone, and deionized (DI) water. A brass mask was employed to print a button circuit measuring 10 cm square onto the glass substrate. The conductivity of the solution was measured on the platinum (Pt) printed electrode using a digital multimeter ‘Fluke 17B Max Digital Multimeter’.

## Results and Discussions

### 3.1. Characterization

#### 3.1.1. Synthesis of MXene and 2D sheets

HF etching method was employed to synthesize MXene (Ti_3_C_2_T_x_) from MAX phase (Ti_3_AlC_2_) and then delaminated the MXene into 2D MXene sheets using DMSO intercalation method^17^. SEM imaging of the various stages clearly demonstrates the successful transformation of the MAX phase into MXene and subsequently into MXene 2D sheets, as illustrated in **Figure 1**. Notably, **Figure 1b** displays the characteristic accordion-like morphology, while **Figure 1c** exhibits a flake-like structure of 2D MXene sheets. EDX elemental mapping provides clear evidence of a significant reduction in the aluminium (Al) content in MXene (**Figure S2**), with the percentage decreasing to approximately 1.5% compared to the MAX phase, which initially had a composition of around 21%. This observation underscores the effectiveness of the HF etching process in efficiently removing aluminium from the MAX phase to form MXene. The UV-Vis spectra of the three phases showing the characteristic absorbance which is in-line with the previously reported data^24^. Hydrodynamic diameter by dynamic light scattering (DLS) shows a significant decrease in the particle size after delamination of the MXene in to 2D sheets (**Figure 1e**). The hydrodynamic diameter of MXene was recorded around 450nm while 2D sheets was around 250 nm. The hydrodynamic diameter measurements revealed that the size of MXene particles was approximately 450nm, whereas the 2D sheets exhibited a smaller diameter of around 250 nm.

**Figure 1.**
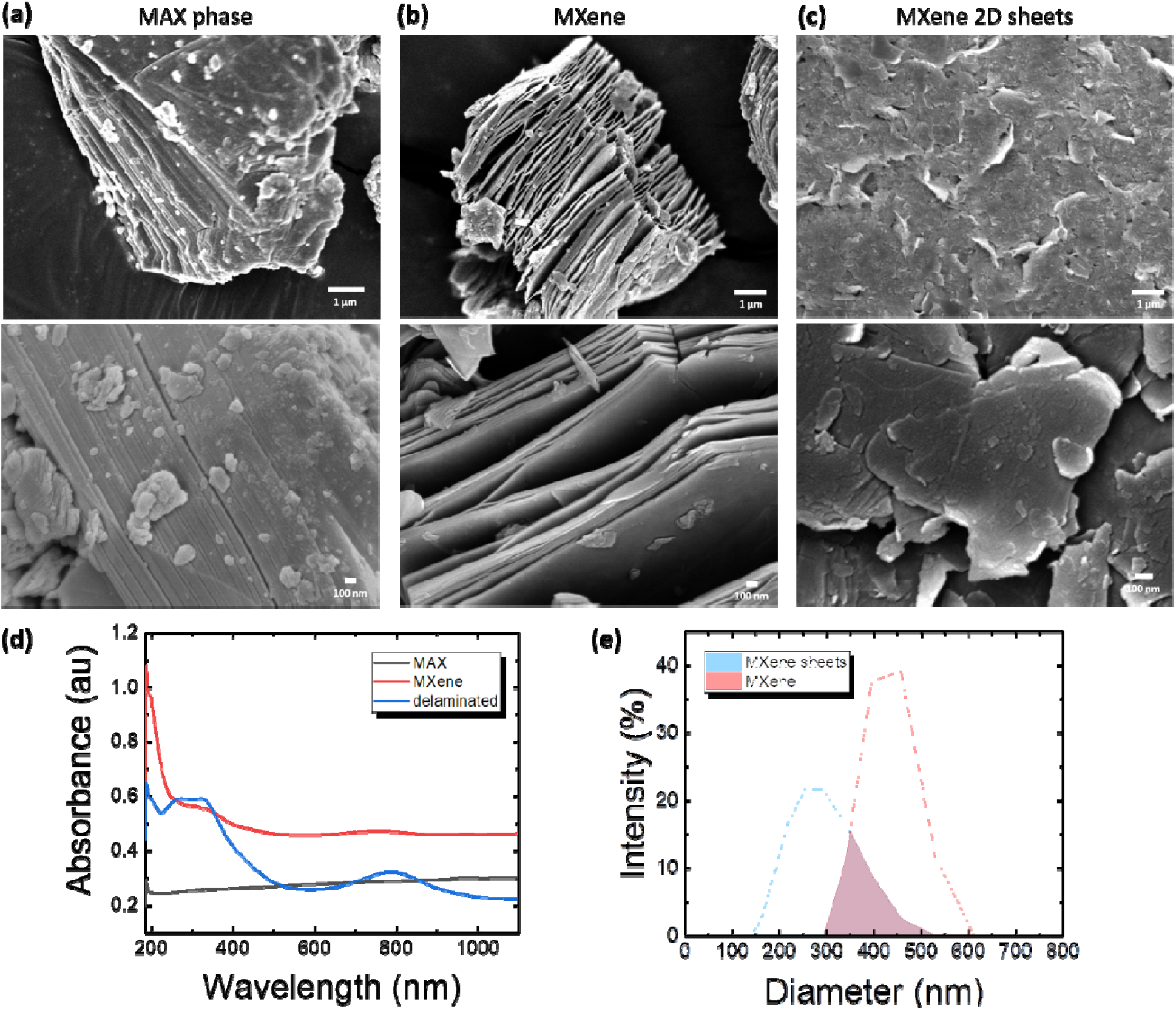
Characterization of MXene formation and it’s delamination into 2D MXene sheets. FE-SEM images of **(a)** MAX phase, **(b)** MXene and, **(c)** delaminated 2D MXene sheets. The upper images are of 10000x zoom and lower images are of higher magnification i.e., 30000x zoom. **(d)** UV-Vis spectra of MAX phase, MXene and 2D MXene sheets. **(e)** Hydrodynamic diameters of the MXene and delaminated 2D MXene sheets by dynamic light scattering.

#### 3.1.2. Aptamer immobilization on 2D MXene sheets

The 2D MXene sheets were amine (-NH_2_) functionalize after APTES treatment, and the DNA aptamer (Apt1) and its complimentary sequence (Apt2) were thiol (-SH) modified at 5’end (**Figure 2a**). The -NH_2_ and -SH groups were covalently bind to each other via Sulfo-MBS linker, resulting into aptamer functionalized 2D MXene (Apt-MX). The absorption of UV light at 260 nm wavelength is notably strong in nucleic acids, primarily attributed to the resonance structure of their purine and pyrimidine bases^25^. The immobilization was characterized by UV-Vis spectroscopy of the Apt-MX showed a significant peak at 260 nm and absent in bare 2D MXene (**Figure 2b**). In the FTIR spectrum of the 2D MXene-DNA hydrogel, distinct vibration peaks emerged, indicating the presence of new molecular interactions in comparison to the FTIR spectrum of pure 2D MXene (**Figure 2c**).

**Figure 2.**
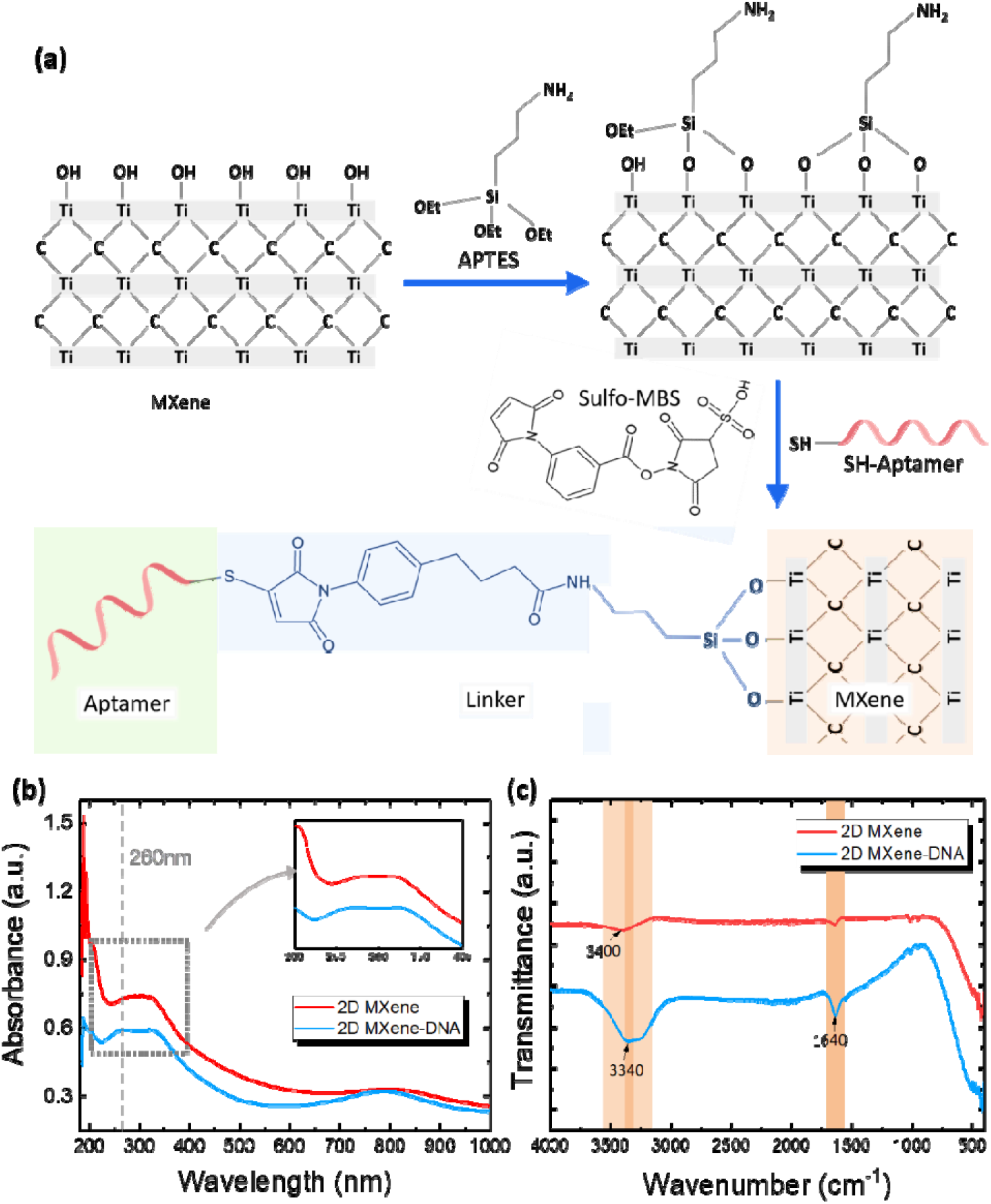
(a) Schematic representation of aptamer immobilization on 2D MXene sheets. Where MXene sheets were first amine functionalized with APTES treatment and then thiol modified aptamer is covalently attached to it by Sulfo-MBS crosslinker. (b) UV-Vis spectroscopy analysis of bare 2D MXene sheets (2D MXene) and aptamer immobilized 2D MXene sheets (2D MXene-DNA).

The broad peak at 3400 cm^-1^ corresponds to the free -OH group, which is noticeably absent or submerged with other peak in 2D MXene-DNA hydrogel, possible due to the APTES binding to the -OH on the surface of 2D MXene^26^. Additionally, an increased signal at 1640 cm^-1^ is evident in the FTIR spectrum of the 2D MXene-DNA hydrogel, attributed to the surface water molecules, including hydroxyl groups in the organic part^27,28^. Furthermore, a new peak emerged at 3340 cm-1 (ranging from 3350 to 3310 cm-1). This spectral addition is indicative of a secondary amine, likely a result of the linkage between the free amine of APTES and Sulfo-MBS^29^. These distinct changes in the FTIR spectrum provide insights into the molecular interactions and compositional alterations occurred in the 2D MXene-DNA hydrogel.

### 3.2. Functionality of thrombin binding aptamer (TBA)

**Figure 3a** shows the mobility disparity between two single-stranded DNA (ssDNA) oligos: Apt1 (21 nucleotides) and Apt2 (21 nucleotides). Both Apt1 and Apt2 have been thiol-modified at their 5’ ends. As Apt2 is a smaller oligo, it exhibits higher mobility compared to Apt1. The bands of both Apt1 and Apt2 appear lighter in shade, which is a result of ethidium bromide (EtBr) staining. EtBr is known to preferentially intercalate with double-stranded DNA (dsDNA), resulting in lighter bands for single-stranded DNA. While the mixture of Apt1 and Apt2, displays even lower mobility and a darker shade, indicating successful hybridization between the two oligos. This observation confirms the clear formation of a hybrid duplex between Apt1 and Apt2, a crucial outcome for their use as a crosslinker.

**Figure 3.**
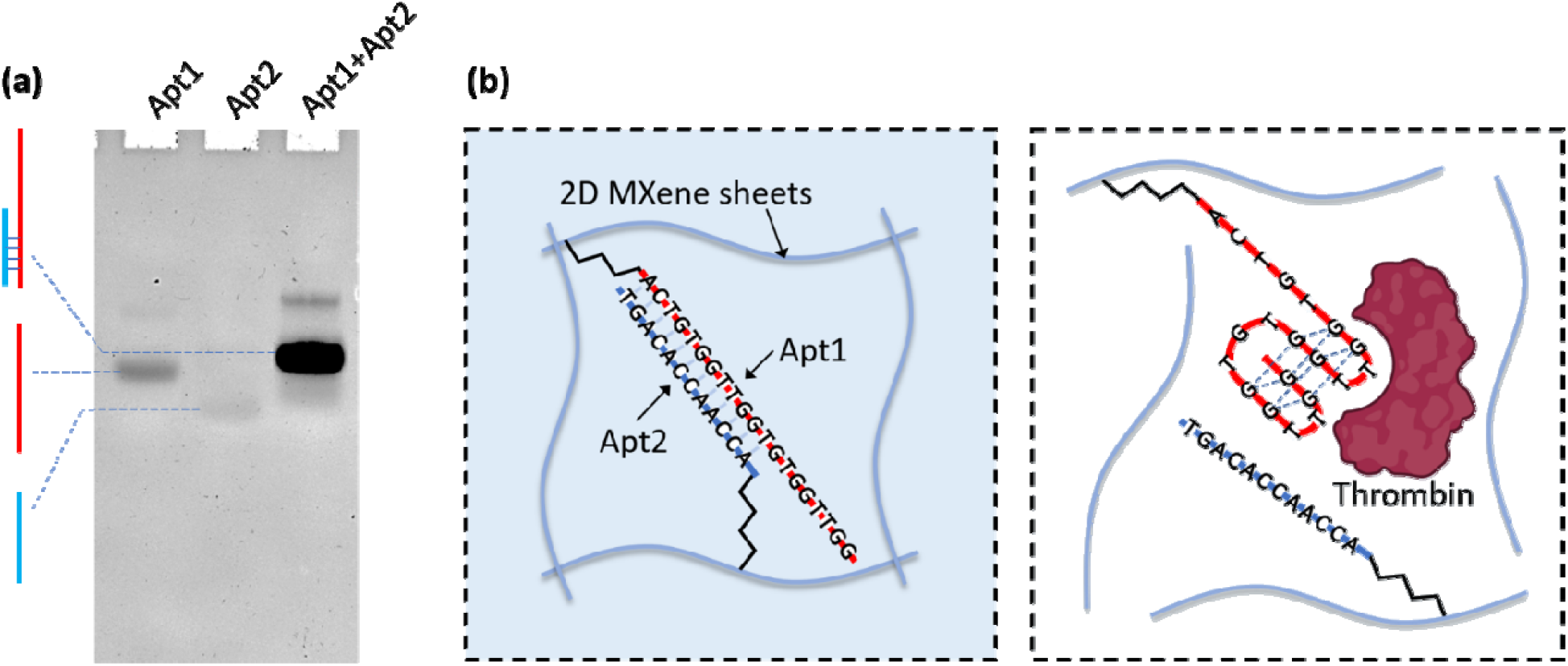
(a) Mobility disparity between the DNA oligos. The Apt2 with lower molecular weight moving faster than the relatively heavier Apt1, while the dsDNA is heaviest with higher EtBr intercalating capability. (b) Schematic representation of working of DNA aptamer oligos (Apt1+Apt2) as crosslinker and keep the 2D MXene sheets together. In presence of thrombin the Apt1 leaves Apt2 and breaks the crosslinking, thus act as an actuator.

The DNA oligos are immobilized on 2D MXene sheets, creating a cohesive structure. In the absence of thrombin, Apt1 (TBA) forms a duplex with Apt2, contributing to the stability of the structure. However, when thrombin is introduced, the binding dynamics change. Apt1 dissociates from Apt2 and undergoes a conformational change, folding into a G-quadruplex structure (**Figure 3b**). This transformation occurs due to the higher affinity of the TBA for thrombin, TBA binds to the fibrinogen-recognition site (exosite), with a dissociation constant (K_d_) of ∼100 nM. Consequently, the effective binding of Apt1 with thrombin leads to the disruption of the Apt1-Apt2 hybrid, causing the structure to become unstable. The formation of the G-quadruplex structure in Apt1 during thrombin binding alters the interactions with Apt2, resulting in the loss of cohesive forces that were initially responsible for maintaining the integrity of the MXene sheets. This instability is a crucial step in the process, as it enables Apt1 to specifically interact with thrombin and fulfil its intended role in detection.

### 3.3. Effect of thrombin on the hybrid MXene-DNA hydrogel

The hybrid hydrogel has been ingeniously designed to achieve strong bonding between the 2D MXene sheets through Watson-Crick complementary base pairing (H-bonding) of the immobilized DNA fragments (Apt1 and Apt2). This unique arrangement results in a densely packed hydrogel where electroconductive MXene sheets remains in close proximity with each other, leading to high current density or lower resistivity. However, when thrombin molecules are introduced, a fascinating change occurs. The presence of thrombin causes Apt1 to break its H-bonding with Apt2 and instead bind with thrombin, forming a G-Quadruplex structure (as shown in **Figure 3b**). This transformation alters the hydrogel’s morphology and properties. The FE-SEM images of the MXene-DNA composition reveal a dense and intricate structure where 2D MXene sheets appear interconnected. On the other hand, in the presence of thrombin, the same composition exhibits a different behaviour, with discrete 2D MXene sheets observed (**Figure 4**). This structural change reflects the specific binding of thrombin with Apt1 and its influence on the overall arrangement of the hybrid hydrogel.

**Figure 4.**
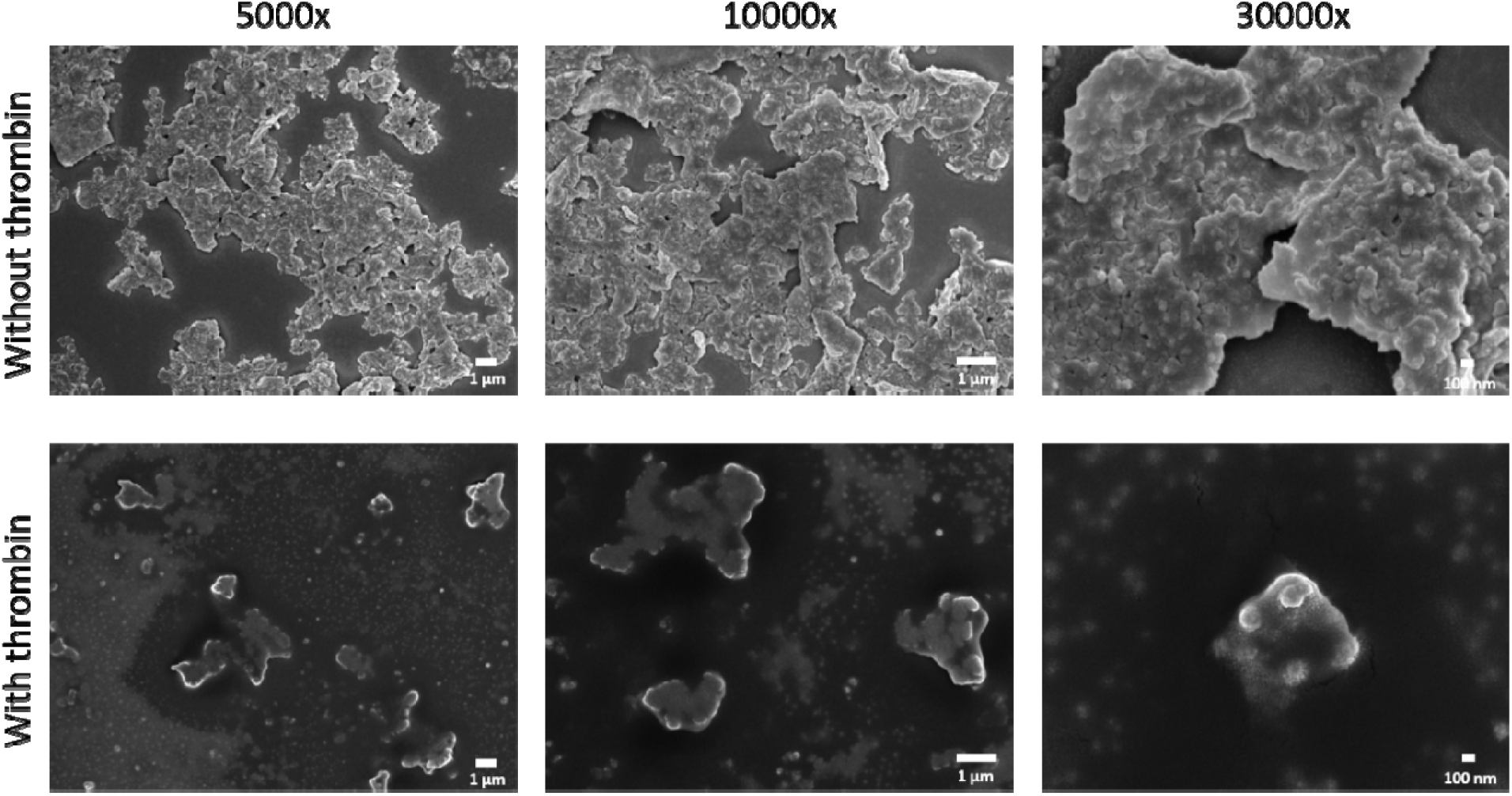
FE-SEM images of the MXene-DNA hydrogel with and without thrombin at different magnification i.e., 5000x, 10000x and 30000x. The upper row shows the hydrogel’s structure before the introduction of thrombin, where 2D MXene sheets exhibit a network-like arrangement. At 30,000x magnification, three or more 2D MXene sheets can be observed in close proximity and interconnected, forming a continuous network. On the other hand, lower row displays the hydrogel after thrombin addition, revealing discrete islands of the 2D MXene sheets. The presence of discrete islands indicates the absence of crosslinking, suggesting a structural change induced by thrombin.

### 3.4. Thrombin level detection

The 2D sheets of MXene are known to possess electrical conductivity when combined with organic matter or polymer. In a study conducted by An et al. in 2018, they demonstrated the electrical conductivity of 2D MXene when integrated with nylon fibres. Specifically, they coated nylon fibres 2D MXene, and with this combination exhibited notable electrical conductivity. Remarkably, the researchers successfully closed the circuit using the MXene-polymer complex, and as a result, a light-emitting diode (LED) bulb illuminated^30^. Researches have shown the capability of MXene-polymer complex as stretchable and bendable conductive polymer to detect the motion or movement in the body^31,32^. It is known from electrochemical impedance spectroscopy data that the 3D structure of MXene hydrogel facilitate better conductivity then the MXene powder^26^.

Here in this study, the developed system intended to measure the blood thrombin levels by isolating the serum from the blood. The thrombin containing blood serum will induce the physical changes while adding it to the MXene-DNA hydrogel, which could be measured by a multimeter in terms of resistance (**Figure 5a**). The physical structural changes in the developed MXene-DNA composite resulted in a change in resistance when introduced with the target molecule, which is thrombin in this case. To facilitate the resistance measurement of the MXene-DNA hydrogel, we employed a Pt printed glass electrode, as shown in **Figure 5b**. Specifically, we used a laser-printed mask made of brass to create a specific button circuit with an area of 1 cm2 (**Figure 5c**). The Pt electrode was applied on glass slide using the sputtering technique to form the button circuit (as shown in **Figure S1**). It is important to note that the printed circuit remains open and does not allow any current flow until a conductive liquid is dropped onto it. This conductive liquid is the MXene-DNA hydrogel containing the blood serum. The presence of thrombin in the serum induces structural changes in the MXene-DNA composite, leading to altered resistance. By measuring the resistance changes in the hydrogel using the Pt printed glass electrode and multimeter setup, we effectively gauge the thrombin levels in the blood serum.

**Figure 5.**
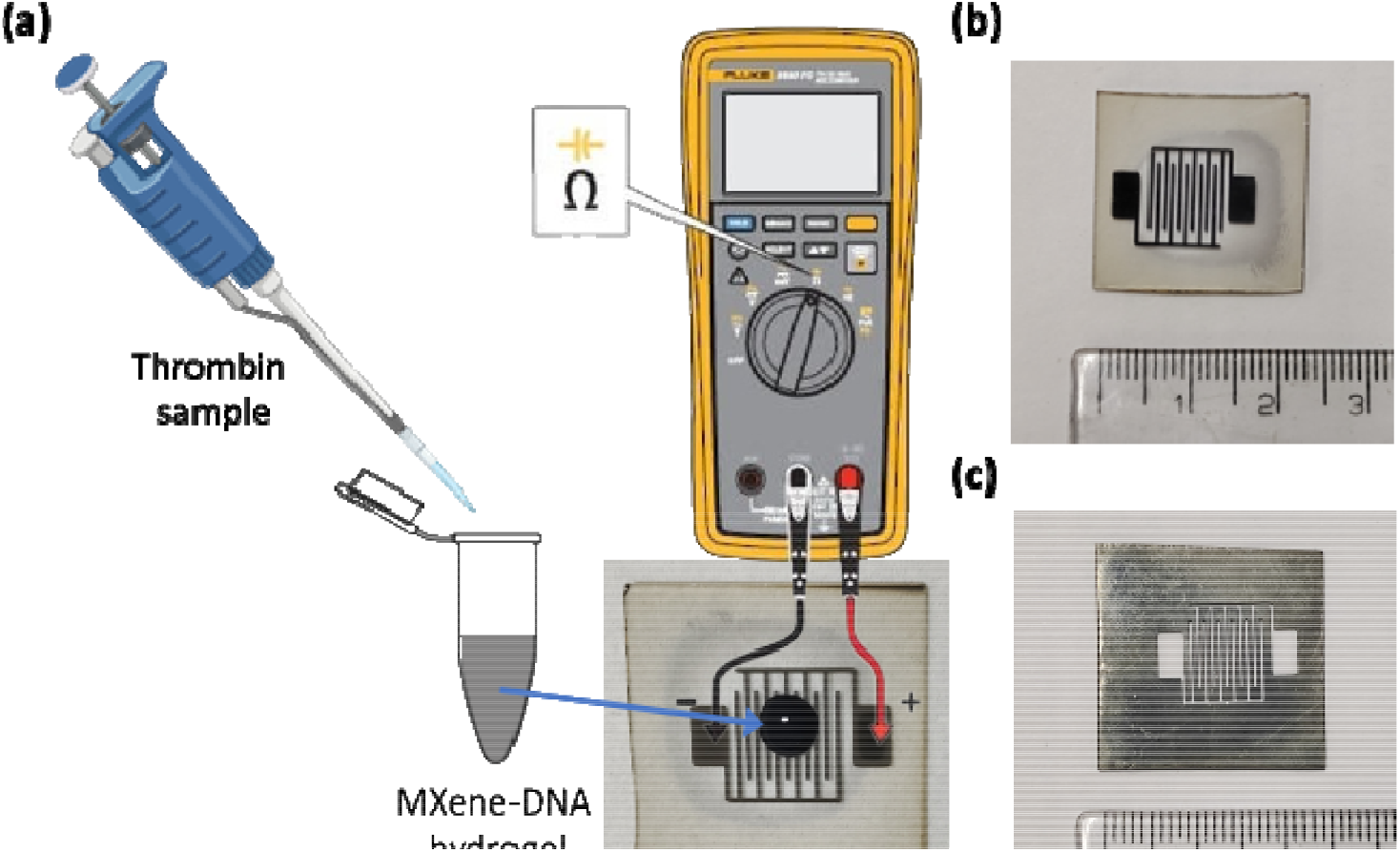
(a) Schematic representation of the workflow of the thrombin detection system. Addition of a sample containing thrombin to the MXene-DNA hydrogel will lead to a specific resistance after applying on to the Pt printed button circuit. (b) Digital photograph of Pt printed button circuit on glass substrate by sputtering technique. (c) Digital photograph of the bras mask used for printing the button circuit on glass substrate. (The scale is in cm)

To determine the thrombin levels in unknown samples, we conducted resistance measurements using four known concentrations and subsequently plotted a calibration curve. In a healthy individual, the normal blood thrombin levels typically range from 50 to 100 mg/L^33^. To establish the calibration curve, we utilized concentrations of 10, 50, 100, and 200 mg/L of human blood thrombin (**Figure 6b**). These known concentrations served as reference points for relating resistance values to thrombin levels in the subsequent measurements of unknown samples. The calibration curve allows us to quantitatively assess the thrombin content in the unknown sample. Upon fitting a linear regression to the four-point calibration data, we obtained an R-squared value (coefficient of determination) of approximately 0.988. Although this value is not considered ideal, it still allows us to obtain reliable primary results. The R-squared value of 0.988 indicates that the linear regression model captures a significant amount of the variation in the data, providing a reasonably good fit for estimating thrombin concentrations within the tested range. Providing a reasonably good fit for estimating thrombin concentrations within the tested range and a limit of detection (LOD) of 0.1698 mg/L and a resolution of 6.51 mg/L. During our experimentation, we analysed an artificial sample with an unknown thrombin concentration of 60 mg/L. Using the calibration curve, we previously constructed, we calculated the thrombin level in the sample to be 68 mg/L. This result indicates a deviation of 8-10% from the expected concentration, providing valuable insights into the accuracy and reliability of our calibration curve for thrombin detection.

**Figure 6.**
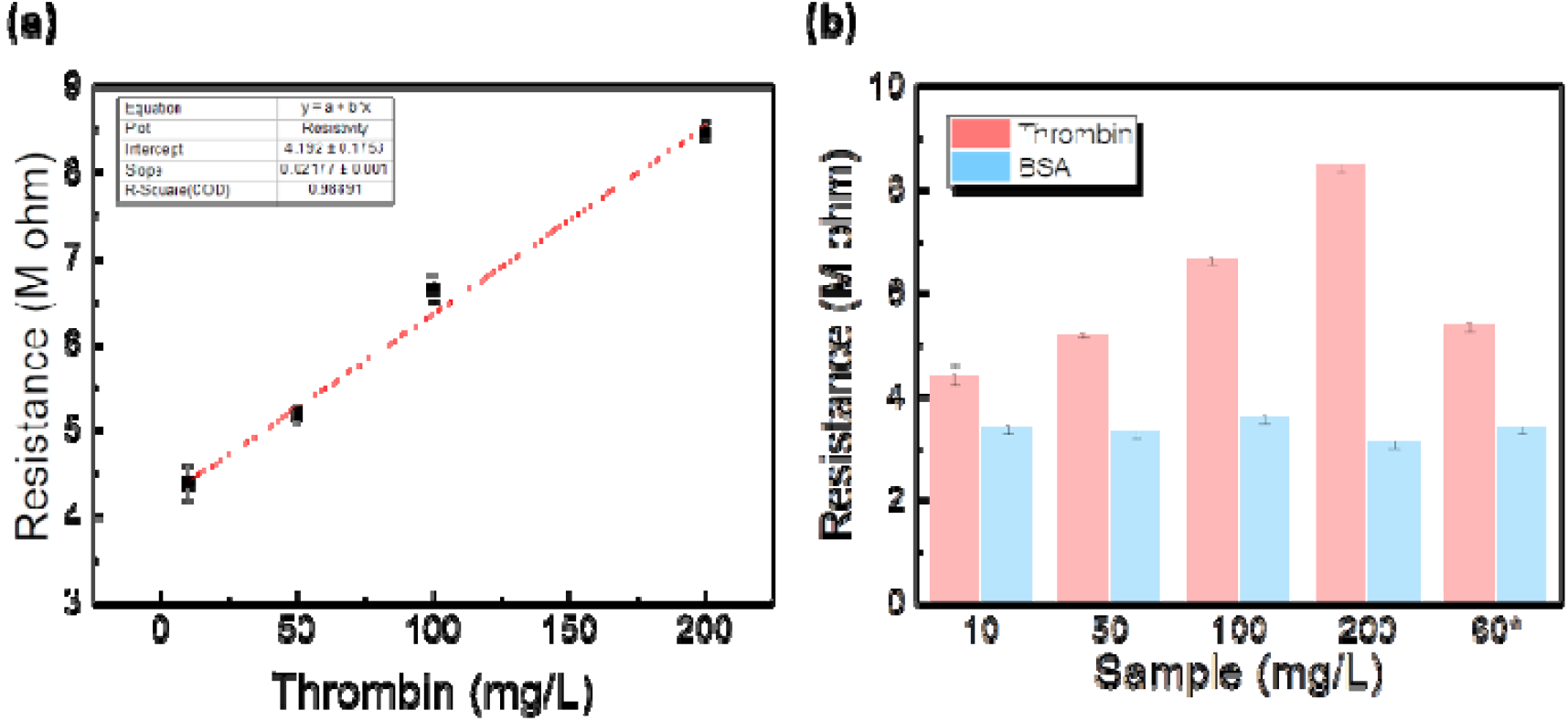
Detection of thrombin concentration. (a) Calibration curve from the different concentrations of thrombin. (Linear regression fit in red line) (b) Bar graph of resistance value observed from different concentration of thrombin and BSA. 60*= artificial sample of 60 mg/L concentration.

Moreover, in order to assess the system’s specificity towards thrombin, we conducted experiments using bovine serum albumin (BSA), a widely recognized model protein. Interestingly, our findings revealed no significant change in the resistance of the 2D MXene DNA hydrogel when exposed to BSA. This outcome serves as a clear indication that the developed system exclusively responds to thrombin, reinforcing its selectivity and specificity towards this target analyte. To demonstrate the robustness and versatility of the approach, a similar hybrid hydrogel was developed for the detection of ampicillin in water samples. Ampicillin is a widely used antibiotic from the penicillin class of medicine, which could be hazardous to human health if inappropriately used^34^. An increase in resistance was observed with increasing concentrations of ampicillin, indicating response to the target molecule with a calculated LOD for ampicillin of 0.1155 mg/L and a resolution of 0.0597 mg/L. Additionally, when tested with tetracycline, the hybrid hydrogel showed specificity for ampicillin over tetracycline (Figure S3).

## Conclusions

In this study, we have demonstrated a novel hybrid hydrogel by harnessing the unique properties of DNA and MXene materials. The DNA aptamer and its partially complementary oligo were successfully immobilized onto 2D MXene sheets, leading to the development of a well-characterized MXene-DNA hybrid hydrogel. We employed various analytical techniques, including UV-Vis spectroscopy, FT-IR spectroscopy, DLS and FE-SEM, to thoroughly characterize the hydrogel’s formation. The DNA fragments acted as crosslinkers, facilitating the binding of 2D MXene sheets in a compact 3D network within the hydrogel. However, upon introducing thrombin, the crosslinkers underwent conformational changes, resulting in a loosening of the hydrogel structure. This physical transformation in the hydrogel was ingeniously utilized to create a thrombin detection system. The specific changes in morphology triggered by the presence of thrombin led to alterations in the resistivity of the MXene-DNA complex. Our developed detection system exhibited the ability to analyze different concentrations of thrombin in artificial samples with an acceptable deviation of 8-10%. The sensitivity of the multimeter used may account for this variation. Furthermore, the system demonstrated specificity towards its target, thrombin, as evidenced by its negligible response to BSA.

Overall, this work establishes a proof-of-concept for a potential electronic device that can precisely sense a target analyte. This system’s versatility shines through the potential to adapt it for virtually any molecule or disease biomarker by modifying the aptamer. The combination of MXene and DNA in the form of a hybrid hydrogel opens up exciting opportunities for various sensing and biomedical applications, where specific conformational changes can be harnessed to develop sensitive and selective detection systems.

## Author Contributions

The manuscript was written through contributions of all authors. All authors have given approval to the final version of the manuscript. CG and VM conceived the idea and planned the experiments. VM designed the experiments, performed all the experiments, and analyzed the data, wrote the first draft of the manuscript. VM, CG, DB analyzed the data and helped with final draft of the manuscript. DB and CG procured the funding for the project.

## Acknowledgements

We sincerely thank the members of CG and DB labs for constant support and discussions. Gratitude is extended to Mr. Akshant Kumawat for his invaluable assistance with FE-SEM imaging.VM thank IITGN, MoE GoI for the PhD fellowships. DB thanks SERB-DST GoI for Ramanujan Fellowship and Core research grant. This work was also funded by Gujcost-DST, GSBTM funds. The central instrumentation facilities (CIF) and Common research and technology development hub (CRTDH) at IITGN are gratefully acknowledged.

## References

(1) Khajouei, S.; Ravan, H.; Ebrahimi, A. DNA Hydrogel-Empowered Biosensing. Advances in Colloid and Interface Science 2020, 275, 102060. 10.1016/j.cis.2019.102060.

(2) Morya, V.; Walia, S.; Mandal, B. B.; Ghoroi, C.; Bhatia, D. Functional DNA Based Hydrogels: Development, Properties and Biological Applications. ACS Biomater. Sci. Eng. 2020, 6 (11), 6021–6035. 10.1021/acsbiomaterials.0c01125.

(3) Hu, Y.; Ying, J. Y. Reconfigurable A-Motif, i-Motif and Triplex Nucleic Acids for Smart pH-Responsive DNA Hydrogels. Materials Today 2023, 63, 188–209. 10.1016/j.mattod.2022.12.003.

(4) Morya, V.; Shukla, A. K.; Ghoroi, C.; Bhatia, D. pH-Responsive and Reversible A-Motif-Based DNA Hydrogel: Synthesis and Biosensing Application**. ChemBioChem 2023, 24 (10), e202300067. 10.1002/cbic.202300067.

(5) Zhao, L.; Li, L.; Yang, G.; Wei, B.; Ma, Y.; Qu, F. Aptamer Functionalized DNA Hydrogels: Design, Applications and Kinetics. Biosensors and Bioelectronics 2021, 194, 113597. 10.1016/j.bios.2021.113597.

(6) Zhou, L.; Jiao, X.; Liu, S.; Hao, M.; Cheng, S.; Zhang, P.; Wen, Y. Functional DNA-Based Hydrogel Intelligent Materials for Biomedical Applications. J. Mater. Chem. B 2020, 8 (10), 1991–2009. 10.1039/C9TB02716E.

(7) Yang, H.; Liu, H.; Kang, H.; Tan, W. Engineering Target-Responsive Hydrogels Based on Aptamer−Target Interactions. J. Am. Chem. Soc. 2008, 130 (20), 6320–6321. 10.1021/ja801339w.

(8) Ning, Y.; Hu, J.; Lu, F. Aptamers Used for Biosensors and Targeted Therapy. Biomedicine & Pharmacotherapy 2020, 132, 110902. 10.1016/j.biopha.2020.110902.

(9) Giannetti, A.; Tombelli, S. Aptamer Optical Switches: From Biosensing to Intracellular Sensing. Sensors and Actuators Reports 2021, 3, 100030. 10.1016/j.snr.2021.100030.

(10) Klug, S. J.; Famulok, M. All You Wanted to Know about SELEX. Mol Biol Rep 1994, 20 (2), 97–107. 10.1007/BF00996358.

(11) Zhou, W.; Huang, P.-J. J.; Ding, J.; Liu, J. Aptamer-Based Biosensors for Biomedical Diagnostics. Analyst 2014, 139 (11), 2627–2640. 10.1039/C4AN00132J.

(12) Wang, C.; Liu, X.; Wulf, V.; Vázquez-González, M.; Fadeev, M.; Willner, I. DNA-Based Hydrogels Loaded with Au Nanoparticles or Au Nanorods: Thermoresponsive Plasmonic Matrices for Shape-Memory, Self-Healing, Controlled Release, and Mechanical Applications. ACS Nano 2019, 13 (3), 3424–3433. 10.1021/acsnano.8b09470.

(13) Zhang, L.; Jean, S. R.; Ahmed, S.; Aldridge, P. M.; Li, X.; Fan, F.; Sargent, E. H.; Kelley, S. O. Multifunctional Quantum Dot DNA Hydrogels. Nat Commun 2017, 8 (1), 381. 10.1038/s41467-017-00298-w.

(14) Basu, S.; Chakraborty, A.; Alkiswani, A.-R. I.; Shamiya, Y.; Paul, A. Investigation of a 2D WS2 Nanosheet-Reinforced Tough DNA Hydrogel as a Biomedical Scaffold: Preparation and in Vitro Characterization. Mater. Adv. 2022, 3 (2), 946–952. 10.1039/D1MA00897H.

(15) Xu, Y.; Wu, Q.; Sun, Y.; Bai, H.; Shi, G. Three-Dimensional Self-Assembly of Graphene Oxide and DNA into Multifunctional Hydrogels. ACS Nano 2010, 4 (12), 7358–7362. 10.1021/nn1027104.

(16) Hosseini, M.; Rahmanian, V.; Pirzada, T.; Frick, N.; Krissanaprasit, A.; Khan, S. A.; LaBean, T. H. DNA Aerogels and DNA-Wrapped CNT Aerogels for Neuromorphic Applications. Materials Today Bio 2022, 16, 100440. 10.1016/j.mtbio.2022.100440.

(17) Alhabeb, M.; Maleski, K.; Anasori, B.; Lelyukh, P.; Clark, L.; Sin, S.; Gogotsi, Y. Guidelines for Synthesis and Processing of Two-Dimensional Titanium Carbide (Ti3C2Tx MXene). Chem. Mater. 2017, 29 (18), 7633–7644. 10.1021/acs.chemmater.7b02847.

(18) Negrier, C.; Shima, M.; Hoffman, M. The Central Role of Thrombin in Bleeding Disorders. Blood Reviews 2019, 38, 100582. 10.1016/j.blre.2019.05.006.

(19) Doherty, T. M.; Kelley, A. Bleeding Disorders. In StatPearls; StatPearls Publishing: Treasure Island (FL), 2023.

(20) Brummel, K. E.; Butenas, S.; Mann, K. G. An Integrated Study of Fibrinogen during Blood Coagulation*. Journal of Biological Chemistry 1999, 274 (32), 22862–22870. 10.1074/jbc.274.32.22862.

(21) Pelzer, H.; Schwarz, A.; Heimburger, N. Determination of Human Thrombin-Antithrombin III Complex in Plasma with an Enzyme-Linked Immunosorbent Assay. Thromb Haemost 1988, 59 (1), 101–106.

(22) Bichler, J.; Heit, J. A.; Owen, W. G. Detection of Thrombin in Human Blood by Ex-Vivo Hirudin. Thrombosis Research 1996, 84 (4), 289–294. 10.1016/S0049-3848(96)00189-2.

(23) Zhang, G.; Wang, T.; Xu, Z.; Liu, M.; Shen, C.; Meng, Q. Synthesis of Amino-Functionalized Ti3C2Tx MXene by Alkalization-Grafting Modification for Efficient Lead Adsorption. Chem. Commun. 2020, 56 (76), 11283–11286. 10.1039/D0CC04265J.

(24) Ji, J.; Zhao, L.; Shen, Y.; Liu, S.; Zhang, Y. Covalent Stabilization and Functionalization of MXene via Silylation Reactions with Improved Surface Properties. FlatChem 2019, 17, 100128. 10.1016/j.flatc.2019.100128.

(25) Olson, N. D.; Morrow, J. B. DNA Extract Characterization Process for Microbial Detection Methods Development and Validation. BMC Res Notes 2012, 5, 668. 10.1186/1756-0500-5-668.

(26) Deng, Y.; Shang, T.; Wu, Z.; Tao, Y.; Luo, C.; Liang, J.; Han, D.; Lyu, R.; Qi, C.; Lv, W.; Kang, F.; Yang, Q.-H. Fast Gelation of Ti3C2Tx MXene Initiated by Metal Ions. Advanced Materials 2019, 31 (43), 1902432. 10.1002/adma.201902432.

(27) Dong, F.; Wang, H.; Wu, Z. One-Step “Green” Synthetic Approach for Mesoporous C-Doped Titanium Dioxide with Efficient Visible Light Photocatalytic Activity. J. Phys. Chem. C 2009, 113 (38), 16717–16723. 10.1021/jp9049654.

(28) Lee, J.; Jo, W. Control of Methyl Tertiary-Butyl Ether via Carbon-Doped Photocatalysts under Visible-Light Irradiation. Environmental Engineering Research 2012, 17. 10.4491/eer.2012.17.4.179.

(29) Smith, B. C. Infrared Spectral Interpretation: A Systematic Approach; CRC Press, 2018.

(30) An, H.; Habib, T.; Shah, S.; Gao, H.; Radovic, M.; Green, M. J.; Lutkenhaus, J. L. Surface-Agnostic Highly Stretchable and Bendable Conductive MXene Multilayers. Science Advances 2018, 4 (3), eaaq0118. 10.1126/sciadv.aaq0118.

(31) Guo, J.; Yu, Y.; Zhang, H.; Sun, L.; Zhao, Y. Elastic MXene Hydrogel Microfiber-Derived Electronic Skin for Joint Monitoring. ACS Appl. Mater. Interfaces 2021, 13 (40), 47800–47806. 10.1021/acsami.1c10311.

(32) Guo, B.; He, S.; Yao, M.; Tan, Z.; Li, X.; Liu, M.; Yu, C.; Liang, L.; Zhao, Z.; Guo, Z.; Shi, M.; Wei, Y.; Zhang, H.; Yao, F.; Li, J. MXene-Containing Anisotropic Hydrogels Strain Sensors with Enhanced Sensing Performance for Human Motion Monitoring and Wireless Transmission. Chemical Engineering Journal 2023, 461, 142099. 10.1016/j.cej.2023.142099.

(33) Putnam, F. W. The Plasma Proteins: Structure, Function, and Genetic Control; Elsevier, 2013.

(34) Yuan, R.; Yan, Z.; Shaga, A.; He, H. Design and Fabrication of an Electrochemical Sensing Platform Based on a Porous Organic Polymer for Ultrasensitive Ampicillin Detection. Sensors and Actuators B: Chemical 2021, 327, 128949. 10.1016/j.snb.2020.128949.

