## Supplementary material for "2D MXene-DNA Hybrid Hydrogel for Thrombin Detection: A Versatile Approach for Biomedical Sensing": Supplimentary file

Table S1. Sequence of oligos used in the study.

| # | Oligo | Sequence (5’-3’) |
| --- | --- | --- |
| 1 | Apt1 (Aptamer for thrombin) | 5′- [ThiC6] -ACT GTG GTT GGT GTG GTT GG-3′ |
| 2 | Apt2 (cDNA for Apt1) | 5’- [ThiC6] -ACC AAC CAC AGT-3’ |
| 3 | Apt3 (Aptamer for Ampicillin) | 5′- [ThiC6] -GCG GGC GGT TGT ATA GCG G-3’ |
| 4 | Apt4 (cDNA for Apt3) | 5′- [ThiC6] -CCG CTA TAC AAC -3' |

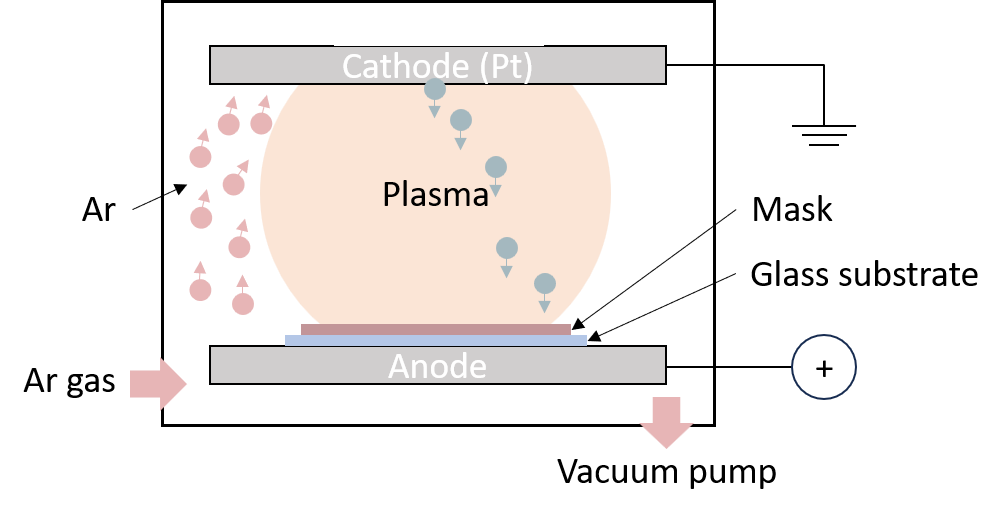

Figure S1. Schematic representation of the setup for sputtering to develop the Pt printed glass electrode with a mask.

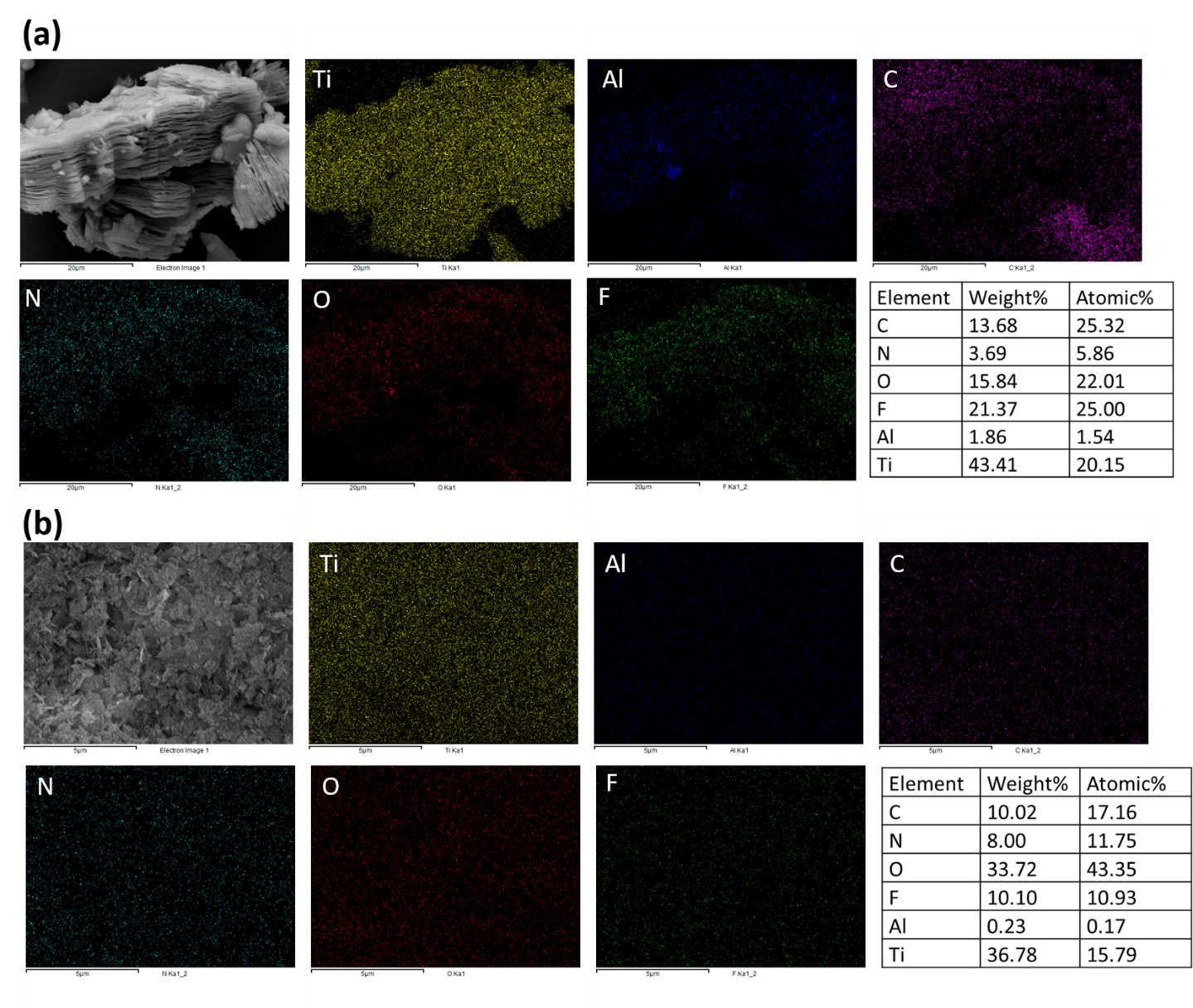

Figure S2. Energy disperive x-ray (EDX) elemental mapping of (a) MXene and (b) MXene-DNA complex.

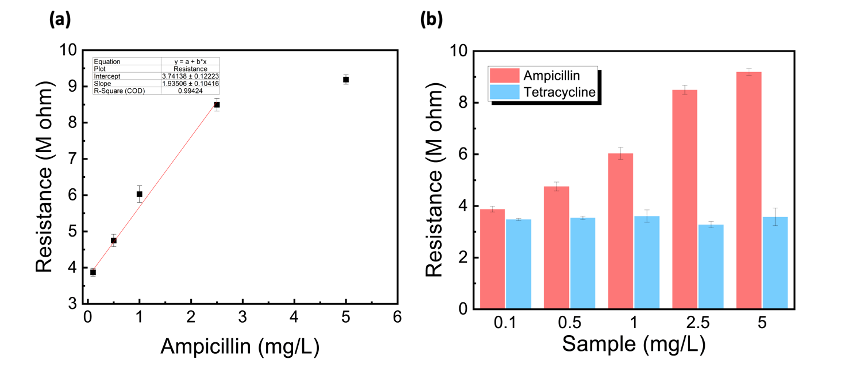

Figure S3. Detection of Ampicillin concentration. (a) Calibration curve from the different concentrations of Ampicillin. (Linear regression fit for intial four points is in red line) (b) Bar graph of resistance value observed from different concentrations of Ampicillin and Tetracycline. The calculated LOD for Ampicillin was 0.1155 mg/L and the resolution was 0.0597 mg/L.

The limit of detection (LOD) and the resolution calculated by the following equations^1^.

$$LOD=mulyiplication factor (3.3) \times SD_{blank}$$

$$SD_{blank}= \sqrt{\frac{\sum{(X_{i}-\bar{X}_{blank})}^{2}}{N_{blank}-1}}$$

Where,

$X_{i}$is each individual measurement of the blank signal.

$\bar{X}_{blank}$is the mean of the blank signal.

$N_{blank}$is the number of measurements in the blank.

$$Resolution=\frac{LOD}{Slope}$$
